# Self-assembling all-enzyme hydrogels for biocatalytic flow processes

**DOI:** 10.1101/240325

**Authors:** Theo Peschke, Sabrina Gallus, Patrick Bitterwolf, Yong Hu, Claude Oelschlaeger, Norbert Willenbacher, Kersten S. Rabe, Christof M. Niemeyer

## Abstract

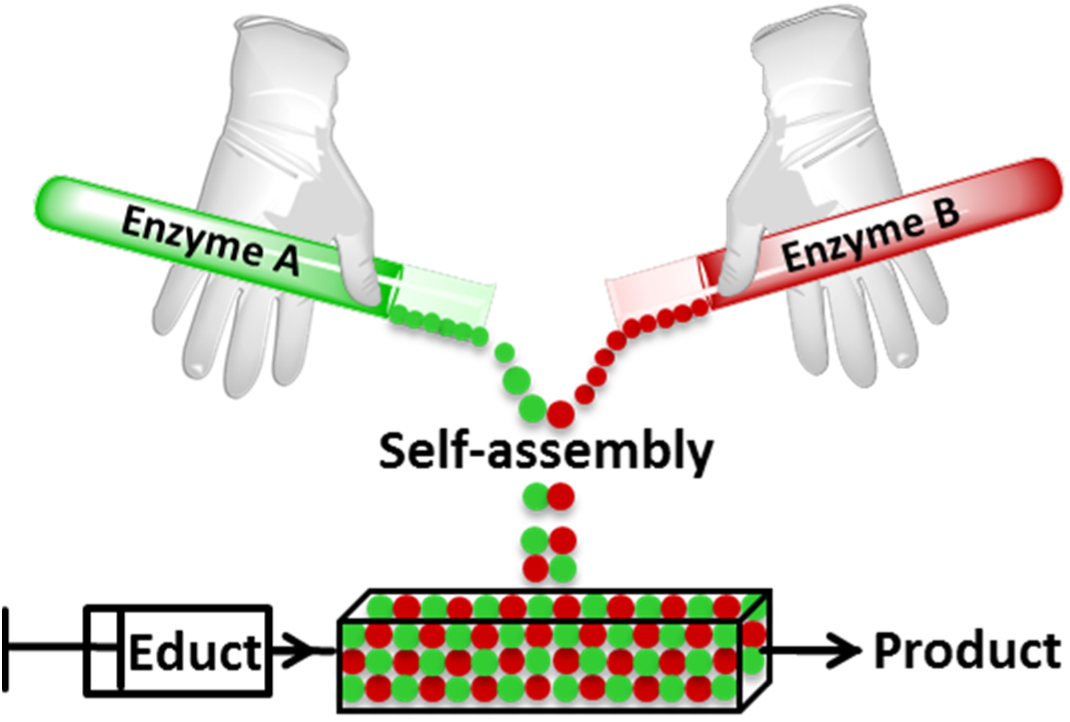

We describe the construction of binary self-assembling all-enzyme hydrogels that are comprised entirely of two tetrameric globular enzymes, the stereoselective dehydrogenase LbADH and the cofactor-regenerating glucose 1-dehydrogenase GDH. The enzymes were genetically fused with a SpyTag or SpyCatcher domain, respectively, to generate two complementary homo-tetrameric building blocks that polymerise under physiological conditions into porous hydrogels. The biocatalytic gels were used for the highly stereoselective reduction of a prochiral diketone substrate where they showed the typical behaviour of the coupled kinetics of coenzyme regenerating reactions in the substrate channelling regime. They effectively sequestrate the NADPH cofactor even under continuous flow conditions. Owing to their sticky nature, the gels can be readily mounted in simple microfluidic reactors without the need for supportive membranes. The reactors revealed extraordinary high space-time yields with nearly quantitative conversion (>95%), excellent stereoselectivity (d.r. > 99:1), and total turnover numbers of the expensive cofactor NADP(H) that are amongst the highest values ever reported.

---

Biocatalysis is a green and sustainable technology that is widely considered as a key domain of industrial (‘white’) biotechnology, which is expected to have an enormous impact on the emergence of biobased economy^1, 2^. Towards the goal of the efficient use of renewable biomass as an alternative to petrochemical synthesis for sustainable production processes and energy supply in the future, multistep enzymatic cascade reactions are currently attracting much attention^3–8^. Such cascades are abundant in living systems, for instance, to maintain the regulation of metabolic activity or signal transduction^9–11^. Their exploitation for technical processes, however, remains difficult because usually compartmentalisation is needed to prevent the multiple reactions from spreading and unproductive crosstalk. This biomimetic strategy has been effectively implemented in conventional organic synthesis by spatially separating sequential transformations into individual reaction vessels that are fluidically coupled with each other. This approach, dubbed as ‘flow chemistry’, is spurred by a high degree of machine-assisted automation^12^ and has yielded impressive synthesis campaigns for small molecules in recent times^13–15^.

The development of biocatalytic flow chemistry-like systems is underway with a special focus on microfluidic bioreactors that offer the generic advantage of a high degree of control over temperature profiles and diffusion-based mixing^8, 16–18^. However, biocatalytic processes are difficult to realize in flow systems because the heterogeneous catalysis regime calls for effective surface immobilization techniques that are more demanding for enzymes than for conventional organo(metallic) catalysts^19^. Common techniques for enzyme immobilization inside microstructured flow channels, such as simple non-specific physisorption or chemical crosslinking, are often hampered by adverse effects on the enzymes’ catalytic activity, whereas (bio)orthogonal one-point immobilization strategies, mediated by genetically encoded immobilization tags, are better suited for the immobilization of delicate enzymes^20^. However, it remains the problem that the amount of immobilized biocatalyst is limited by the effective surface area. To overcome this limitation, pseudo-3D interfacial layers comprised of synthetic polymers or micro-/nanoparticles can be used to increase the number of binding sites and, thus, the loading capacity for enzymes^21, 22^. Since this approach requires additional coupling steps with potential drawbacks for biocatalytic activity, *in situ* generation of pure enzyme polymeric networks would provide an ideal solution for the loading of microfluidic reactors with large amounts of active biocatalysts. Hydrogels are porous polymers that can be constructed from natural or synthetic structural proteins^23, 24^. A recently established protein gelation strategy utilizes a pair of genetically encoded reactive partners, SpyTag and SpyCatcher, that spontaneously form covalent isopeptide linkages under physiological conditions^25–29^. While these protein hydrogels are being explored for applications in biomedical sciences, such as cell encapsulation and tissue engineering, strategies for their exploitation in biocatalysis remain underdeveloped.

We here present the first example of self-assembling all-enzyme hydrogels that display extraordinary high space-time yields in biocatalytic flow processes and show the behaviour of coupled kinetics of coenzyme regenerating reactions in the substrate channelling regime. Specifically, we constructed a self-ligating binary enzyme polymer using two tetrameric globular enzymes, the stereoselective dehydrogenase LbADH and the cofactor-regenerating glucose 1-dehydrogenase GDH. The two enzymes were genetically fused with a SpyTag or SpyCatcher, respectively, thereby generating the two complementary homo-tetrameric building blocks of the polymer. Gelation under physiological conditions led to spontaneous formation of porous hydrogels that displayed the characteristic activity of the pure enzymes in the highly stereoselective reduction of a prochiral diketone substrate. Importantly, the gels were capable to effectively sequestrate the NADP(H) cofactor even under continuous flow conditions. Owing to their sticky nature, the gels are readily mounted into microfluidic reactors without the need for supportive membranes or other means for catalyst immobilization. The reactors revealed extraordinary high space-time yields with nearly quantitative conversion (>95%), excellent stereoselectivity (d.r. > 99:1), and total turnover numbers of the expensive cofactor NADP(H) that are amongst the highest values ever reported.

## Results

### Construction of all-enzyme hydrogels

We choose two widely used homotetrameric enzymes, the highly (*R*)-selective alcohol dehydrogenase LbADH (EC 1.1.1.2) from *Lactobacillus brevis* and the nicotinamide adenine dinucleotide phosphate (NADPH)-regenerating glucose 1-dehydrogenase GDH (EC 1.1.1.47) from *Bacillus subtilis*. Both enzymes were genetically fused with either the SpyTag (ST) or the SpyCatcher (SC) in addition to a hexahistidin (His) tag tethered to the same terminus of the protein (Figure 1A). The ST/SC system enables the rapid crosslinking of the two tetravalent protein building blocks through the formation of covalent isopeptide bonds under physiological conditions^30^. The proteins were overexpressed in *E. coli* and purified to homogeneity by Ni-NTA affinity chromatography (Supplementary Figure 1). Initial electrophoretic analysis of enzyme gelation confirmed that polymerisation only occurs when both enzymes bear the complementary binding sites (Supplementary Figure 2). A more detailed investigation of the polymerisation reaction was accomplished by dynamic light scattering (DLS) analysis (Figure 1B, C). As expected, the time-dependent formation of protein clusters occurred on time scales of minutes to hours, depending on the concentration of the two enzyme building blocks (Figure 1B). In homogeneous solution particles with average size of up to 65 nm were formed that further fused to a viscous liquid and even free-standing hydrogel piece upon further desiccation of the solvent (Figure 1A, B). Variation of the stoichiometric ratio of the two enzyme building blocks showed the fastest increase of the hydrodynamic diameter at equimolar ratio (Figure 1C). Analysis of the gel’s morphology by scanning electron microscopy (SEM) and atomic force microscopy (AFM) revealed no clearly distinctive ultrastructure, however, particle-like features were evident in both SEM and AFM images (Figure 1A, Supplementary Figure 3).

**Figure 1.**
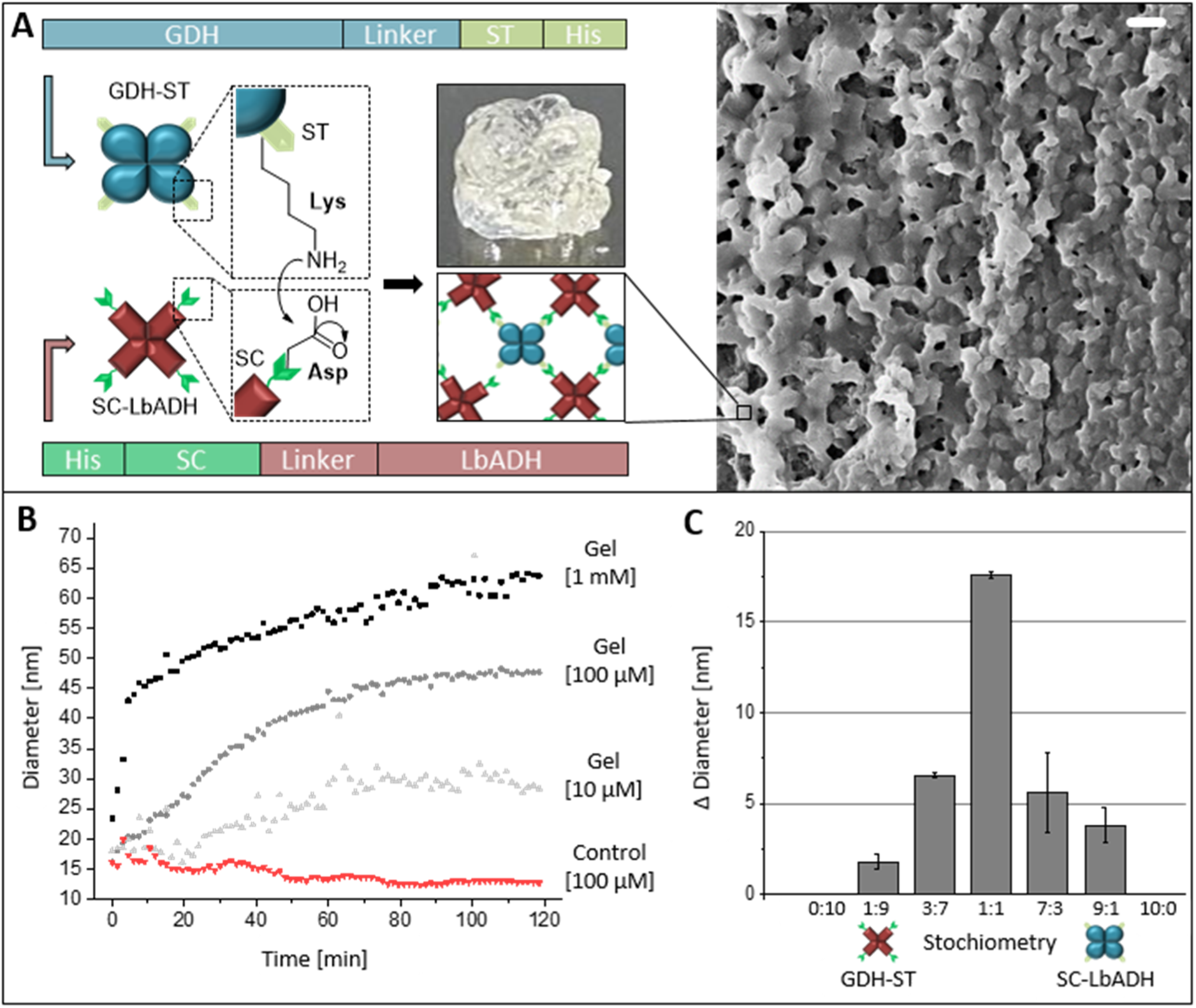
Design, formation and morphological characterization of the self-assembled all-enzyme hydrogels. (**A**) Schematic illustration of the two homotetrameric enzyme building blocks, GDH-ST and SC-LbADH, that can self-assemble to a hydrogel via formation of covalent isopeptide bonds. Photograph and representative SEM image of the hydrogel; scalebar 300 nm; for additional morphological characterization see Supplementary Figure 3. (**B**) Time-and concentration-dependent increase in hydrodynamic diameter (Z-average), determined at 25°C by DLS. Total protein concentrations are given; the control contained equimolar amounts of GDH-ST and LbADH lacking the SC domain. (**C**) Stoichiometry-dependent increase in particle diameter observed in the initial 30 min after mixing of the two enzyme building blocks (100 µM total subunit concentration, 25 °C).

The observation of distinctively sized particle populations suggests that multiple alternating layers of the enzymes are formed upon covalent linkage until a critical size regime is reached where further growth is determined by the reversible interactions between the subunits of the tetrameric GDH and LbADH enzymes. As compared to chemically cross-linked polymers, the gelation of the enzyme hydrogels occurs rather slowly, similar as observed for synthetic elastin hydrogels, where the polymerising proteins first form discrete particles that merge to form larger spheres and further coalesce into open linked networks of a porous hydrogel network^31^. To further elucidate their material properties, the enzyme hydrogels were analysed by optical microrheology based on multiple particle tracking (MPT) analysis^32^ (Supplementary Figure 4). The method revealed that the hydrogel has a homogeneous structure on the micrometer length scale with a *G*_0_ = 20±7 Pa, an average mesh size ξ = 60±7 nm and a pore size <200 nm that is in the range of typical microfiltration membranes^33, 34^. Based on these data, we hypothesize that the GDH-ST/SC-LbADH hydrogels have a hierarchical structure and dynamic morphology that is determined by the reversible interactions between the enzyme’s subunits (see also Supplementary Figure 5).

### Biocatalytic properties of GDH-ST/SC-LbADH hydrogels

Owing to its relevance for stereochemistry and natural product synthesis^35^, we chose the prochiral C_S_-symmetrical 5-nitrononane-2,8-dione (NDK) **1 (**Figure 2) as the substrate for benchmarking the biocatalytic activity of the all-enzyme hydrogels. Depending on the stereoselectivity of a given ketoreductase, either one or both carbonyl groups of NDK are reduced to form diastereomeric hydroxyketones **2** or diols **3**, respectively, and all products can be readily quantified by chiral HPLC analysis^35^ (Supplementary Figures 6, 7). We had previously established that particle-immobilized LbADH converts NDK with very high stereoselectivity into (*R*)-*syn/anti*-hydroxyketones **2c/d** (e.r.>99:1; d.r. ~60:40), which are further reduced to form the (*R,R*)-configured pseudo *C_2_*-diol **3d**^36^. We now used the NDK reaction to initially profile the SC-LbADH and GDH-ST building blocks, which revealed a slightly decreased (30%) and increased (22%) specific activity, respectively, as compared to the untagged enzymes (Supplementary Table S1).

**Figure 2.**
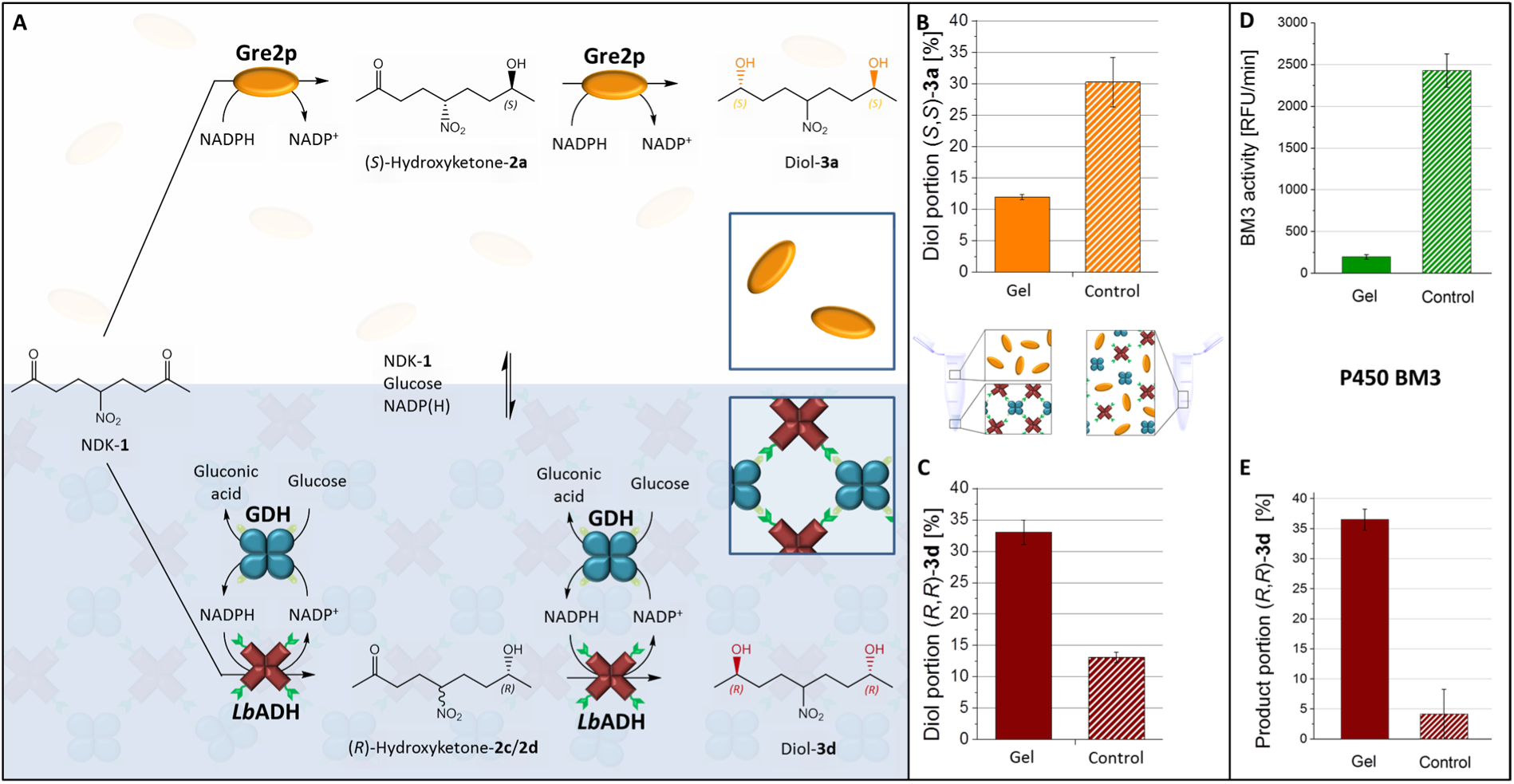
Compartmentalised competition assay using GDH-ST/SC-LbADH hydrogels and Gre2p or P450 BM3 in the bulk solution as competitor for NADP(H). Controls were performed with non-assembling enzymes and substrates, all mixed in homogeneous solution. (**A**) Reaction scheme of the compartmentalised competition assay where NDK-**1** is reduced by either the (*S*)-selective Gre2p (orange) in the liquid bulk phase or the (*R*)-selective LbADH (red) in the gel phase that also contains the NADPH-regenerating GDH (blue). The control contained GDH lacking the ST domain and LbADH-SC (**B**) The bars show the amounts of (*S,S*)-**3a** diol portion produced by Gre2p or (**C**) (*R,R*)-**3d** diol produced by LbADH. (**D, E**) Formation of fluorescent BM3 product and (*R,R*)-**3d** diol, respectively, in the competition assay, where P450 BM3 in the liquid bulk phase competes for NADP(H). Note that in the heterogeneous two-phase competition (gel) about 7-fold lower or higher amounts of BM3 product (**D**) or (*R,R*)-**3d** diol product (**E**) are formed, respectively, as compared to the controls with unassembled enzymes in homogeneous solution. For further details and concentration-dependent product distributions, see Supplementary Figures 9, 10). Error bars indicate the standard deviation, obtained from at least two independent experiments.

The kinetics of NDK reduction was then compared between the GDH-ST/SC-LbADH hydrogel and a control of the unassembled mixed proteins. To this end, hydrogels were prepared into which NADP^+^ was included during polymerisation and solvent evaporation. After swelling of the hydrogel, reaction buffer, containing NDK and glucose, was added and product formation was monitored by chiral HPLC analysis (Supplementary Figure 8). We found that the total activity of the unassembled enzyme system was nearly three-fold higher than that of the hydrogel and we attribute this observation to a restricted diffusion inside the hydrogel. However, the hydrogel was rapidly forming the (*R*,*R*)-**3d** diol with only small amounts of the intermediate hydroxyketones, whereas the unassembled enzymes produced almost exclusively the hydroxyketones. These results suggested that the gel functions as a compartment wherein the highly concentrated and tightly packed enzymes lead to retainment of hydroxyketone intermediates and their favored direct reduction to the (*R*,*R*)-**3d** diol.

It is evident from natural examples that compartmentalized reaction systems offer advantages when intermediate species are prone to escape into competing reaction channels^3–8^. The GDH-ST/SC-LbADH hydrogels can be used to investigate this phenomenon because the NADPH consuming LbADH reduction is spatially and kinetically coupled to the NADPH regeneration by GDH. To explore whether escape of the intermediate species (NADP(H) and hydroxyketones) from the GDH-ST/SC-LbADH hydrogels is affected by a competing reaction, we set up a two-phase system wherein polymerised enzyme hydrogel with entrapped NADP^+^ was covered with a solution containing another trackable NADPH-consuming enzyme, the (*S*)-selective methylglyoxal reductase Gre2p (EC 1.1.1.283) from *Saccharomyces cerevisiae* (Figure 2A). The substrates for the two-step reaction, NDK and glucose, were provided only in the fluid phase that contained the competing Gre2p. Analysis of NDK conversion by chiral HPLC allowed to quantitatively determine the amounts of product formed by either the (*S*)-selective Gre2p inside the fluid phase or the (*R*)-selective LbADH in the gel phase. Control experiments were performed with non-assembling enzymes and substrates, all mixed in homogeneous solution.

We observed that formation of the (*S,S*)- and (*R,R*)-configured diol products was shifted about three-fold, from 30% to 12% (*S,S*)-**3a** and 14% to 33% (*R,R*)-**3d** diol portion, respectively, when the competition experiment was carried out in the two-phase system instead of the homogeneous solution (Figure 2B, 2C, respectively). Furthermore, this change in product distribution was not substantially altered by the concentration of the competing Gre2p enzyme (Supplementary Figure 9). Likewise, a similar compartmentalised competition assay, carried out with NADPH-consuming P450 BM3 as competing enzyme (Figure 2D), revealed that the product portion of (*R,R*)-**3d** diol was increased even seven fold upon compartmentalised co-localization of LbADH and GDH inside the hydrogel (Figure 2E). Indeed, the gel-enclosed LbADH was even more active than free enzymes when the competing enzyme was present in high concentrations (Supplementary Figures 9, 10E). We also confirmed that the data from the compartmentalised competition assays are not an artefact, which may originate from the slight differences in specific activity of the tagged and untagged enzymes (Supplementary Figure 11). Altogether, the results confirmed that the hydrogel - with its high concentration of co-localized NADPH-consuming and -regenerating enzymes - indeed functions as an effective reaction compartment that reduces the escape of intermediate reaction species.

To elucidate the influence of the starting conditions with respect to the localisation of the NADP^+^ cofactor, we used the BM3 competition assay. We compared the enzymatic activity depending on whether NADP^+^ was localised in the gel, the liquid bulk phase solution or equally distributed between the two phases at the beginning of the reaction (Supplementary Figure 12). We found that formation of (*R,R*)-**3d** diol was highest in the case of gel-entrapped NADP^+^. However, the productivity of LbADH was reduced only by about 20% when NADP^+^ was localised in the bulk solution at the reaction start. This finding not only emphasised the effectiveness of cofactor entrapment but it also suggested that NADP(H) was readily sequestrated in the gel phase. We reasoned that both aspects would be advantageous for applications in biocatalytic flow processes, where is a high demand for strategies for cofactor minimisation and easy applicable carrier-free enzyme immobilisation^2^.

To investigate the process stability of the GDH-ST/SC-LbADH hydrogels under flow conditions, we used a PDMS chip with a flow channel of 150 µl volume that was completely filled with the all-enzyme hydrogel (Figure 3A-E). In initial experiments, the channel was perfused with reaction buffer containing NADP^+^, glucose and NDK at a flowrate of 10 µl/min. As expected, the hydrogel effectively retained the immobilised enzymes and produced the (*R,R*)-**3d** diol for prolonged times, while unassembled enzyme mixtures were rapidly washed out of the reactor (Figure 3F, see also Supplementary Figure 13). Importantly, the fluidic reaction control enabled an almost quantitative conversion of NDK to (*R,R*)-**3d** diol. Determination of the flow rate-dependency of the formation of (*R,R*)-**3d** diol product and corresponding space-time yields (STY, Figure 3G) revealed that the reactor could be operated at 50-fold higher flowrates and still performed with a 50-fold higher time yield and a more then 3-fold higher STY, than a previously reported packed-bed microreactor that contained bead-immobilized GDH and LbADH^36^.

**Figure 3.**
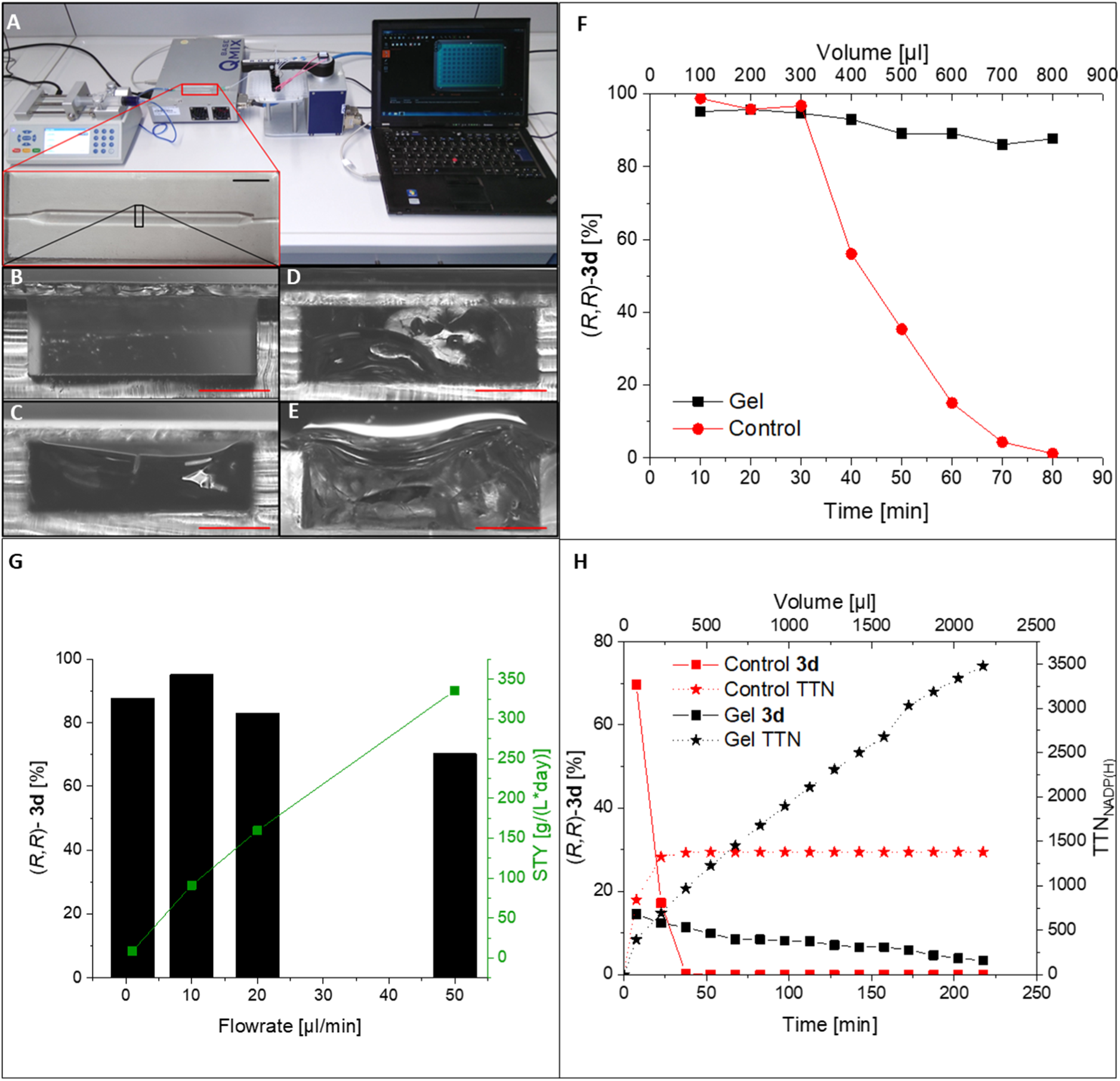
Stereoselective conversion of NDK-1 using a microfluidic reactor loaded with GDH-ST/SC-LbADH hydrogels. (**A**) Overview of the fluidic setup with a magnified image of the chip (scale bar 1 cm). (**B-E**) Zoomed-in cross sections (scale bars 1 mm) of an empty channel (**B**), filled with dried hydrogel (**C**), filled with swollen hydrogel and sealed with a glass slide (**D**) or filled with the swollen hydrogel without sealing (**E**). (**F**) Time dependent production of (*R,R*)-**3d** diol, as determined in the outflow of the enzyme-loaded reactors perfused with continuous cofactor NADP^+^ supply at a flowrate of 10 µl/min. (**G**) Flow rate dependency of the formation of (*R,R*)-**3d** diol product, indicated by the black bars. The corresponding space-time yields (STY) are indicated by the green coloured graph. (**H**) Time dependent production of (*R,R*)-**3d** diol using a reactor that was loaded with GDH-ST/SC-LbADH hydrogel bearing co-entrapped NADP^+^ and perfused with reaction media lacking the NADP^+^ cofactor (flowrate = 10 µl/min). The black graphs represent the amount of (*R,R*)-**3d** diol (squares) and Total Turnover Numbers of NADP(H) (TTN_NADP(H)_, stars) obtained from the polymerised hydrogels, whereas the red graphs indicate the data obtained from a control reactor loaded with unassembled enzymes. Note that the formation of (*R,R*)-**3d** requires two turnovers of NADP(H).

To test whether the all-enzyme hydrogels can be utilised for minimisation of cofactor consumption, microreactors were loaded with hydrogels bearing co-entrapped NADP^+^ and perfused with reaction buffer containing only glucose and NDK (Figure 3H). Indeed, diol product was formed for up to 4 hours indicating the effective retainment of the entrapped cofactor inside the hydrogel over 16 reactor column volumes. The obtained total turnover number of NADP(H) (TTN_NADP(H)_) was clearly in the economically feasible range^37^ and even more then 3-fold higher than those recently reported for a self-sufficient heterogeneous biocatalyst, based on bead-bound ketoreductases with electrostatically co-immobilized NAD(P)H^38, 39^. To the best of our knowledge, the TTN_NADP(H)_ of >3400 observed here is the highest value ever reported for flow processes in devices lacking supportive membranes.

## Discussion

We established a novel class of self-assembled all-enzyme hydrogels that are convenient to prepare and readily mounted in fluidic microreactors. Conventional (multi)enzyme processes require carrier materials, such as beads or membranes, which inevitably ‘dilute’ the specific activity of a given device and, thus, lead to lower space-time yields than those available with all-enzyme systems^2^. Our approach is based on recombinant protein technology, thereby enabling the ‘green’ sustainable production of biocatalytic devices with high catalyst and volume productivity, high stability and low production costs owing to the exclusion of additional expensive carrier materials that require additional efforts for production and disposal.

The ultimately high concentrations of the biocatalytic entities realized in our hydrogels are comparably only to the so-called “cross-linked enzyme aggregates” (CLEA) that can be produced from two or more different proteins in a non-directional fashion by glutaraldehyde mediated unselective cross-linking^40^ or by sophisticated exploitation of metal coordination interactions^41^. These approaches, however, have their limitations in terms of insufficient control over enzyme stoichiometry or sensitivity to environmental conditions (e.g., pH and ion-strength of reaction media), respectively. In contrast, our method is based on robust covalent linkage that is controlled by the versatile SpyTag/SpyCatcher crosslinking technology and even allows for programming of stoichiometries other than the equimolar ratio shown here^29^.

The here described all-enzyme hydrogels reveal the typical behaviour of the coupled kinetics of coenzyme regenerating reactions in the substrate channelling regime. Zhang and Hess have recently emphasized that the rational design of high-efficiency enzyme cascades will benefit not only from spatial proximity of cooperating enzymes but also from balanced stoichiometry^42^ and Castelana et al.^43^ have already demonstrated that enzyme clustering accelerates the processing of intermediates through metabolic channeling. This type of compartmentalisation is manifested in the sequestration of the NADP(H) cofactor and hydroxyketone intermediates inside our all-enzyme hydrogels, which occurs even under continuous flow conditions. In addition to cofactor-regeneration systems, as demonstrated herein, this gel-based compartmentalisation should be particularly useful in biocatalytic multistep reactions where instable or cross reactive intermediates are involved.

Despite excellent enantioselectivity, yields and productivities in the asymmetric reduction of a large variety of carbonyl compounds, the industrial implementation of ketoreductases is often hampered by their dependency on very expensive cofactors, especially when it comes to flow processes where, for instance, NADPH needs to be supplemented continuously^44^. Conventional approaches address this issue by use of ultra-and nanofiltration membrane technology or specifically modified surfaces that minimize or prevent cofactor loss through electrostatic attraction or even covalent immobilisation. While these approaches have led to increased TTN_NADP(H)_ values, they are cost intensive and increase the complexity of production processes, thereby leading to limited economic viability^38, 39, 45, 46^. Our self-assembly approach, in contrast, is straight-forward, scalable and, owing to the gel’s intrinsic material properties, can be readily implemented in arbitrary reactor systems. We therefore believe that the here reported all-enzyme hydrogels will help to improve existing and also devise entirely new strategies for technical biocatalytic systems but possibly also for cell-based production processes, where gel compartments could be harnessed to prevent adverse effects of toxic intermediates.

## Methods

### Preparation of hydrogels

Protein solutions of either GDH-ST-His/His-SC-LbADH (for hydrogel preparation), GDH-His/His-SC-LbADH (soluble enzymes, Control A) or GDH-ST-His/His-LbADH (soluble enzymes, Control B) were diluted in KPi-Mg (100 mM KP_i_ pH 7.5, 1 mM MgCl_2_) to a final concentration of 500 µM in 20 µl for kinetic measurements or 100µl for AFM or SEM sample preparation. Polymerisation of the mixtures was carried out for 1 h at 30°C, 1000 rpm in a thermoshaker. Subsequently, the buffer was evaporated in a 0.2 ml reaction tube with an open lid for 15–17 h at 30°C and under constant centrifugation at 2200 g. For MPT analysis, the samples were supplemented with 0.2 mg/ml ‘dragon’ green fluorescent polystyrene microspheres (200nm diameter; Bangs Laboratories, USA). For experiments with gel entrapped NADP+, the gel solution was supplemented with 10 µM NADP+.

### Determination of enzymatic activity

The specific activities of the enzymes were determined as previously described.^36^ In brief, enzyme assays were performed with a reaction mixture containing 5 mM NDK-**1**, 100 mM glucose, 1 mM NADP^+^, in KP_i_-Mg containing 0,5 µM either His-LbADH, His-SC-LbADH or Gre2p-His as ketoreductase and an excess of 10 µM GDH-His for NADPH-regeneration. The GDH containing reaction mixture was preincubated for at least 30 min at 30°C, before the ketoreductase was added. For the determination of the average specific activity, samples for HPLC analysis were taken after 20 min. Enzyme kinetic reactions were carried out over at least 5 h, samples were taken manually at various time points and were subsequently analysed by chiral HPLC. In order to measure the GDH activity, 0.5 µM GDH-His or GDH-ST-His were incubated together with an excess of 10 µM His-LbADH, using the same conditions as described for the ketoreductases above. For HPLC analysis, 50 µL of the crude reaction mixtures were extracted with 150 µL ethyl acetate, centrifuged for phase separation, and 75 µl of the organic phase were transferred into HPLC vials and evaporated (Concentrator plus, Eppendorf). Values for conversion, enantiomeric-and diastereomeric excess were calculated based on the ratios of HPLC signals detected at 210 nm, as previously described.^35^

### Enzyme kinetics of hydrogel

The dried hydrogels were swollen by addition of 20 µl KP_i_-Mg for 10 minutes under continuous shaking at 25°C, 1000 rpm. Subsequently, a reaction mixture in KP_i_-Mg was added to the swollen hydrogels to a final volume of 200 µl (final concentrations were 1 µM NADP^+^, 5 mM NDK-**1**, 100 mM glucose, 50 µM GDH-ST-His and 50 µM of His-SC-LbADH) and incubated at 30°C, 500 rpm. The same setup was used with GDH-His instead of GDH-ST-His to generate the controls containing unassembled enzymes. The reaction was carried out over at least 4 h at 30°C, 500 rpm and time dependent samples for HPLC analysis were taken manually.

### Compartmentalised competitive assay with Gre2p

Initially, 20 µl KP_i_-Mg were added to the NADP^+^ encapsulating hydrogel or dried enzyme pellets, respectively. After 10 min incubation at 25°C, 1000 rpm, the competition experiment was started by adding 180 µl KP_i_-Mg buffered reaction mixture to obtain final concentrations of 100 mM glucose, 5 mM NDK-**1**, and 0–250 µM Gre2p-His. The reaction mixture was incubated for 26 h at 30°C, 700 rpm and samples were taken manually for chiral HPLC analysis.

### Compartmentalised competitive assay with P450 BM3

The assay was performed similar to the competitive Gre2p assay, except that the P450 BM3 (A74G, F87V)-His mutant was used instead of Gre2p-His and the reaction mixture was additionally supplemented with 100 µM of the BM3 substrate 12-(4-trifluoromethylcoumarin-7-yloxy)dodecanoic acid (TCD)-**4**^47^. The reaction was carried out in 0.2 ml reaction tubes and analysed in a fluorescence plate reader (Synergy MX, BioTek Instruments GmbH). The P450 BM3 (A74G, F87V) catalyses O-dealkylation of TCD-**4** to yield the fluorescent product 7-hydroxy-4-trifluoromethylcoumarin (HTC)-**5**, which was monitored over 2 h (λex = 420 nm, λem = 500 nm). Subsequently the samples were extracted and analyzed by chiral HPLC.

### Flow system analysis of the hydrogels

The PDMS-chips were placed inside an incubator 1000 (Heidolph, Germany, set to 30°C), filled with 150 µl 1000 µM protein solution and incubated for 30 min. This process was repeated 3 times and the PDMS chips were then sealed with a glass slide. The NADP^+^-encapsulating hydrogels contained the final concentration of 10µM NADP^+^. A pumping unit (Fusion 100, Chemyx Inc., USA) with two independent syringe modules equipped with 5 or 20 ml Omnifix syringes (B. Braun Melsungen AG, Germany) was connected to the chip for perfusion of reaction media. The syringes were filled with 5 mL substrate solution containing 5 mM NDK-**1**, 100 mM glucose in KPi-Mg, supplemented with 0.01 % (v/v) sodium azid to avoid fouling and 0 or 1 mM NADP^+^ depending on the individual experiment. The chip outflow was connected to the Compact Positioning System rotAXYS (CETONI, Germany) to allow for automatic fractioning into 96-well plates, previously loaded with 50 µl 7 M CH5N3·HCl to stop all enzymatic reactions. The positioning system was connected to a CETONI neMESYS Base module, which was controlled by the QmixElements-Software. The chip was connected to syringes and the fraction collector by tubings (Inlets: silicone Tygon tubing R3603, ID = 1.6 mm, Saint-Gobain, France; outlets: conventional PTFE tubing, ID = 0.5 mm) using standard cannulas and luer lock fittings.

Further details on the cloning, expression, purification and characterization of the recombinant proteins as well as the chip fabrication, chiral HPLC, DLS, AFM, SEM and MPT microrheological analyses are provided in the Supplementary Information

## Acknowledgments

This work was supported by the Helmholtz programme “BioInterfaces in Technology and Medicine” and DFG project Ni399/15–1. We thank Hatice Mutlu and Arnold Leidner for help with the DLS measurements, Silla Hansen for help with the microfluidic system, Anke Dech, Jens Bauer, Ishtiaq Ahmed, Leonie Hacker for experimental help with protein expression and NDK synthesis, Pia Schwitters and Teresa Burgahn for providing the P450 BM3 plasmid, and Volker Zibat for performing SEM measurements.

## Author Contributions

T.P., S.G. and K.S.R. designed recombinant proteins, developed and characterized the enzyme hydrogels. T.P., S.G., P.B. and Y.H. conducted the SEM analysis. C.O. and N.W. designed and conceived the MPT analysis. T.P. and P.B. performed the flow reactor experiments. T.P., K.S.R. and C.M.N. wrote the manuscript. C.M.N planned and supervised the project. All authors discussed the results and commented on the manuscript.

